# How often are male mosquitoes attracted to humans?

**DOI:** 10.1101/2023.03.08.531798

**Authors:** Véronique Paris, Christopher Hardy, Ary A. Hoffmann, Perran A. Ross

## Abstract

Many mosquito species live close to humans where females feed on human blood. While male mosquitoes do not feed on blood, it has long been recognized that males of some species can be attracted to human hosts. To investigate the frequency of male mosquito attraction to humans, we conducted a literature review and human-baited field trials, as well as laboratory experiments involving males and females of three common Aedes species. Our literature review indicated that male attraction to humans is limited to a small number of species, including Ae. aegypti and Ae. albopictus. In our human-baited field collections, only 4 out of 13 species captured included males. In laboratory experiments, we found that male Ae. notoscriptus and Ae. vigilax showed no attraction to humans, while male Ae. aegypti exhibited persistent attraction for up to 30 minutes. Both male and female Ae. aegypti displayed similar preferences for different human subjects, suggesting that male Ae. aegypti respond to similar cues as females. Additionally, we found that mosquito repellents applied to human skin effectively repelled male mosquitoes. These findings shed light on mosquito behaviour and have implications for mosquito control programs, particularly those involving the release or monitoring of the male mosquito population.

## 1. Introduction

Many insect species exhibit distinct behavioural differences between sexes, often as adaptations in behaviour such as mating, foraging, territoriality and feeding that affect the relative contribution of the sexes to their offspring [1–3]. For example, mate acquisition behaviour in male insects is typically associated with territoriality [4], lekking displays [5], and locating sites where females emerge [6]. Females, on the other hand, more rarely actively search for mates but may focus on accepting males following courtship and nuptial gifts [7–9].

In blood-sucking insects like mosquitoes, males feed on nectar while most females require blood to reproduce and thus have behavioural adaptations for host-seeking and blood-feeding [10,11]. The evolution of blood-feeding in insects is believed to have occurred through different routes, such as accidental biting of vertebrates by plant-sucking insects, which then developed the ability to digest and utilize protein-rich blood [12]. Another possibility is that blood-feeding evolved through the close association between chewing insects and vertebrates, where insects became accustomed to recognizing and biting vertebrates [13]. As blood became crucial for these insects, parallel evolution occurred between insects and their hosts, with the insect developing preference for specific hosts based on cues that optimize reproduction [14]. Anthropophilic mosquitoes exhibit a strong drive to seek out human hosts for blood-feeding and use a combination of cues to locate their target at different spatial scales [15,16]. Once CO_2_ indicates the presence of a human on a broad spatial scale, host cues such as heat and odours are used detect the host once in closer proximity. Mosquitoes feeding on non-human animals also use habitat cues like fresh animal faeces [17]. While CO_2_ is generally considered a host cue [18], there is considerable evidence that it primarily functions as a habitat cue by indicating the general area inhabited by potential hosts [19].

The study of host-seeking behaviour in mosquitoes has traditionally focused on females, as they are the primary vectors of disease transmission. How male mosquitoes recognize habitat and hosts cues remain understudied. However, as the utilization of Wolbachia-infected [20] or sterilized males [21] as a control strategy for reducing mosquito populations becomes increasingly prevalent, understanding the behaviour of male mosquitoes is of growing importance. This is because the efficacy of these methods hinges upon the ability of released males to locate and reproduce with wild females. Male mosquitoes have sophisticated auditory and olfactory systems [11] used to locate females [22,23], nectar and other sugar sources [24], and conspecific males [25]. Despite their inability to blood feed, field observations report that males of Aedes aegypti [26–29] and Ae. albopictus [30] are attracted to humans, with males swarming around and landing on humans. Capture rates of males in both species also increase when traps are baited with CO_2_ or human odour mimics [31–33]. Amos et al [34] confirmed the attraction of Ae. aegypti to humans experimentally under semi-field conditions. In contrast, studies on other mosquito species frequently report no attraction of males to humans and traps that use human cues. For example, studies on Ae. notoscriptus have reported exceptionally low capture rate of males through CO_2_-baited BG traps [35,36], indicating that there may be differences in male behaviour between mosquito species. These may be due to species differences in mating strategies and/or sensory abilities, although the inability to detect male attraction in some cases may be a consequence of study designs which fail to detect male attraction [34].

Species differences in male attraction to humans provide a basis for further investigations into the underlying mechanisms governing this behaviour and how they vary across mosquito species. Species differences are also of applied importance as releases of incompatible or sterile male mosquitoes start to be used to suppress mosquito species; public acceptance of this strategy may be problematic if males are attracted to humans. To investigate species differences, we conducted a literature review of previous observations from field collections that employ human-baited methods, and we present results of our own human-baited field collections of both male and female mosquitoes from various regions in Australia. We also evaluated the attraction of male and female mosquitoes of three common Aedes species to human hosts in laboratory experiments. For species that exhibited human attraction by males, we determined whether preferences for specific human hosts are similar for males and females and tested the effectiveness of mosquito repellents on male mosquitoes.

## 2. Methods

### 2.1. Literature review

We compiled observations on male mosquito attraction to humans from studies on human-baited field collections. We looked for studies that presented catches of both males and females across any mosquito species. We searched the terms “human landing catch mosquito male” as well as “human bait mosquito male” on the Google Scholar platform on September 16, 2022. We went through the first 600 results for each search term to identify references and then also searched for relevant references within the articles. We then searched the Web of Science platform on June 12, 2023, and went through all 492 results for “human landing catch mosquito” and 60 results for “human bait mosquito male”. Studies retained needed to identify mosquitoes to the species level, present numbers of caught mosquitoes for both sexes and present results separated by capturing technique. A study must also have collected mosquitoes through Human Landing Catches (HLC), Human Baited Traps (HBT) or Human Baited Collections (HBC) under field conditions (Figure S1). If these criteria were met, we extracted data on the location of the study, capture method, mosquito species and the number of each sex collected from the text, figures and tables within the article as well as supplementary material. If studies included any interventions or other treatments (e.g., repellents, insecticides, non-human baits), we took care to only extract numbers of catches from control and baseline sites.

### 2.2. Human-baited field trials

We performed human-baited trials in Victoria (VIC), New South Wales (NSW), Australian Capital Territory (ACT), South Australia (SA) and Queensland (QLD) Australia. Detailed information about the location and year of the collections can be found in Table S5 and Figure S2. We ran a total of 115 trails from 2014 – 2022 with 13 different participants (5 female, 8 male; aged 21 – 60) collecting mosquitoes for 0.5 to 1 hr duration at a private residence or public space at any time of the day. Participants were sitting on a chair or bench, exposing both legs from the knee downwards. Mosquitoes were collected when landing or hovering around exposed skin, using mechanical aspirators (Spider & Insect Vac, Select IP Australia Pty Ltd, n = 21), electric rackets (Pestill USB Rechageable Mosqutio & Fly Swatter, Kogan Australia Pty Ltd, n = 23), or tube collection (n = 71). Keys from Dobrorwsky (1965) were used to morphologically identify the species and sex of collected mosquitoes.

Mosquitoes that could not be confidently identified to species level were excluded from the study.

### 2.3. Aedes laboratory experiments

#### 2.3.1. Mosquito strains and maintenance

Laboratory colonies were established from field collections from Cairns, Australia in 2019 (Ae. aegypti) and Brisbane, Australia in 2014 (Ae. notoscriptus) or 2020 (Ae. vigilax). Aedes aegypti and Ae. notoscriptus were reared at 26°C and a 12:12 cycle with a 1 hr dawn and dusk period. Adults were maintained in 30 x 30 x 30 cm BugDorm-1 cages and provided with 70% sucrose solution and females were blood fed using human volunteers (ethics approval from The University of Melbourne 0723847). We collected and partially dried eggs, before hatching them in 3 L of Reverse Osmosis (RO) water containing a total of 0.2 g baker’s yeast. Mosquito larvae were reared on fish food (TetraMin Tropical Fish Food, Tetra, Melle, Germany) and pupae allowed to emerge into cages. Aedes vigilax were reared under identical conditions but adults were maintained in a BugDorm® M4590 Insect rearing cage (93 x 44 x 32 cm) and larvae were reared in 30% saltwater solution (API Aquarium salt, USA).

#### 2.3.2. Male attraction to humans – Aedes species comparison

We conducted experiments on mosquito attraction to humans using three species: Ae. aegypti, Ae. notoscriptus, and Ae. vigilax. The experiments were conducted in a 3 x 3 m tent under constant light levels and at room temperature. Each trial involved releasing 100 males, aged between 1 and 2 weeks, that had previously been allowed to mate, into the tent. The males were given 30 minutes to acclimate before the experiment began. The experiments were filmed using GoPro Hero 10 cameras placed at either end of the tent, with white panels (84.1 x 118.9 cm) as a background. In each trial, one side was baited with a human subject, while the other side was left unbaited as a control. Subjects stood facing the camera with their bare feet and shins in the field of view, with this position remaining consistent across trials. Subjects did not wear any perfume. The side of the baited and unbaited treatment was alternated for each trial. The number of trials, human subjects, and number of days the experiments are summarized in Table S1. The same batch of males was used for multiple trials on the same day but replaced daily. Treatments were recorded for 30 mins using the time-lapse function immediately after the human subject assumed their position inside the tent. The number of mosquitoes in view of the camera was scored every 20 s, distinguishing between males that were in flight and males that landed on the human subject. For data analysis, we calculated the average number of male mosquitoes in each category over the entire trial period.

#### 2.3.3. Mosquito preferences for different human subjects

In our experiments we found that Ae. aegypti males exhibit preferences towards certain human subjects (see Figure S3). While previous research has demonstrated differential attraction of female Ae. aegypti mosquitoes to different human hosts [16,37,38], this has not yet been quantitatively reported in males. We conducted additional experiments in which we used a consistent set of five human subjects (coded A-E) who stood in pairs in opposite positions in the tent setup described in 2.3.2. The subjects were filmed for 5 minutes on each side before the sides were swapped and the procedure was repeated. This was done for each pairwise combination of the five subjects (20 combinations in total), with a fresh batch of males being used for each day of four separate days. The footage was scored as described in 2.3.2. For data analysis, we calculated the average number of male mosquitoes in view (combining flight and landed) over the 5 minutes of each trial for each human subject.

We then tested all pairwise combinations of the same five human subjects for their attraction to female Ae. aegypti and Ae. notoscriptus. We used a two-port olfactometer (30 x 30 x 30 cm) similar to those used in previous studies by Ross et al [39] and Amos et al [34]. The mosquitoes used in this experiment were 6-7 days post-emergence and had been allowed to mate prior to the experiment. We released approximately 50 females into the set-up and allowed them to acclimatise for approximately one minute. A box fan placed at the opposite end of the cage drew air through two traps into the cage. Pairs of subjects placed one hand each in front of one of the traps. After 5 minutes, we closed the entrance to the traps and counted the number of females in each trap and individuals remaining in the cage. The combinations of subjects and sides were randomized until all 20 pairwise comparisons between subjects and sides were completed. We repeated the experiment using the same 5 subjects for another four days using a fresh batch of females each day for a total of 10 replicates (5 per side). Aedes vigilax females were not assessed in this experiment due to relatively low rates of attraction to humans observed in a pilot trial using this olfactometer design.

Prior to data analysis, we calculated a preference index for each person to reflect the relative attraction of each subject by dividing the number of mosquitoes attracted to one human subject by the number of mosquitoes attracted to both subjects. We determined the average preference index for all replicates of each human subject. Statistical analyses were performed using the preference index averaged across replicates.

#### 2.3.4. Effect of mosquito repellent on male mosquitoes that show attraction to humans

After we confirmed that male Ae. aegypti show attraction to humans in our tent experiments, we tested whether they are repelled by a commercial mosquito repellent (Aerogard tropical strength insect repellent, Reckitt Benckiser, NSW, Australia) containing 191 g/kg Diethyltoluamide and 40 g/kg N-Octyl Bicycloheptene Dicarboximide. We used the same tent setup as described in 2.3.2. The repellent was applied to the knees downwards to one of the two human subjects positioned on either site of the tent within 5 min before the trial began. The number of males in view was recorded every 20 s for 10 minutes. The person wearing the repellent and the sides of the treatment and control were randomized. We ran 20 trials over 5 days with a rotation of 9 human subjects, with the batch of 100 Ae. aegypti males replaced each day. The footage was scored as described in 2.3.2. For data analysis, we calculated the average number of male mosquitoes in view (combining flight and landed) over the 5 minutes of each trial.

### 2.4. Data analysis

All statistical analyses were conducted using R (v. 4.1.2) [40]. Wilcoxon-signed-rank tests were used to assess differences in male attraction between three Aedes species. The influence of human subject on the number of male and female Ae. aegypti and female Ae. notoscriptus attracted to humans were assessed by first calculating a preference index for each person to reflect their relative attraction. This involved dividing the number of mosquitoes attracted to one human subject by the number of mosquitoes attracted to both subjects, which was then averaged across the replicates. We then performed an ANOVA, followed by Tukey’s post-hoc tests using this averaged index to test for differences between human subjects. To validate the results obtained through the preference indices, we also build Generalized Linear Mixed-Effects Models using the original data, including the replicate number as a random factor, followed by Tukey’s post-hoc tests. Using the preference indexes (without averaging), we applied Jonckheere-Terpstra tests to determine whether mosquito attraction to one subject was affected by the attractiveness of the other human subject used in a pairwise comparison (ranked apriori by their overall attractiveness). We also ran Mantel tests to compare the matrices of preferences obtained with different groups of mosquitoes to assess whether patterns of preferences differed between species and sexes. Finally, we used a t-test to determine whether the application of mosquito repellent significantly reduced the attraction of male Ae. aegypti to humans.

## 3. Results

### 3.1. Literature review

Our literature review identified 50 studies containing evidence of male mosquito attraction to humans across species using human-baited field collections. A further 355 studies did not meet all our inclusion criteria (Figure S1), including 179 studies that were excluded because they did not specify the sex of the collected mosquitoes.

In the 50 studies involving 137 different mosquito species meeting the inclusion criteria, male catches were reported for 17 species. Among these, only five species (Ae. aegypti, Ae. albopictus, Ae. flavipennis, Ae. riversi and Cx. quinquefasciatus) reported greater than 10% male catches. The evidence for male attraction to humans by Ae. aegypti and Ae. albopictus was robust, with male catches recorded in 20(out of 21) and 17 (out of 19) studies respectively (Figure 1, Table 1).

**Figure 1.**
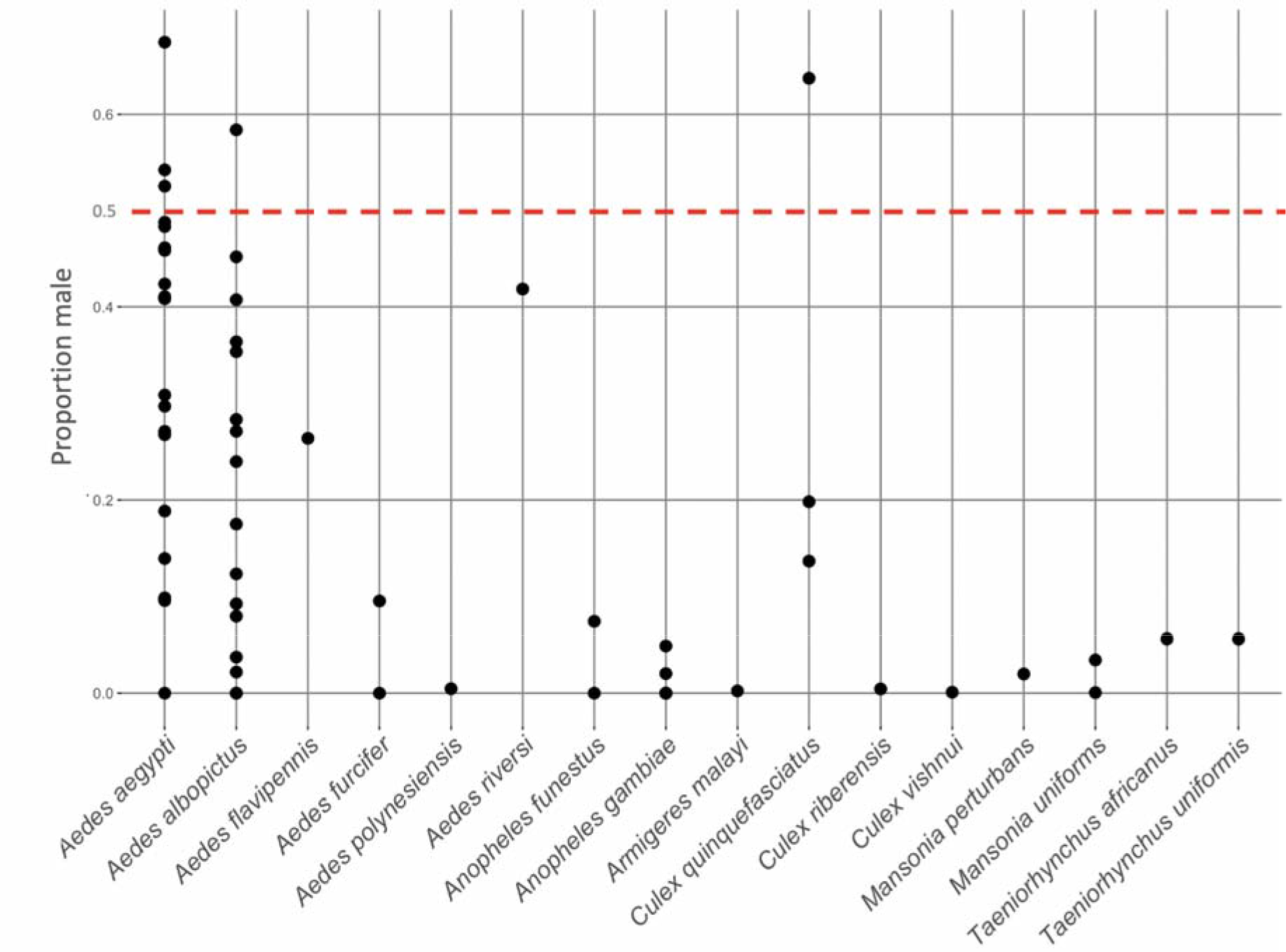
Proportion of males collected across mosquito species from the literature review of human-baited field collections. Dots show the proportion of males collected out of the total catch, with each dot representing a single study. The red dashed line indicates an equal ratio between male and female catches (0.5 proportion). Data are only presented for species with catches having n > 50 individuals and where males were collected. See table S4 for the complete dataset.

**Table 1.**
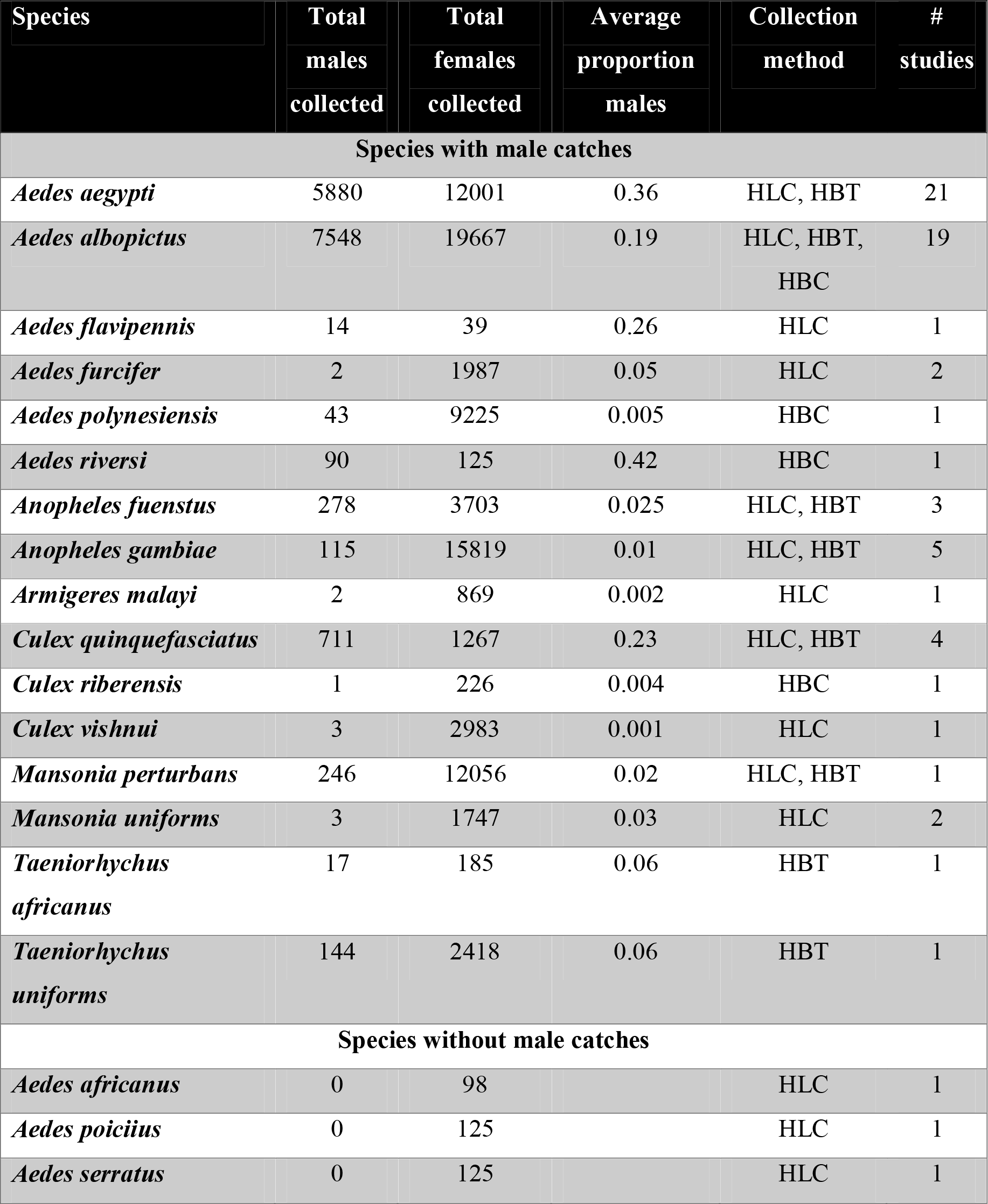
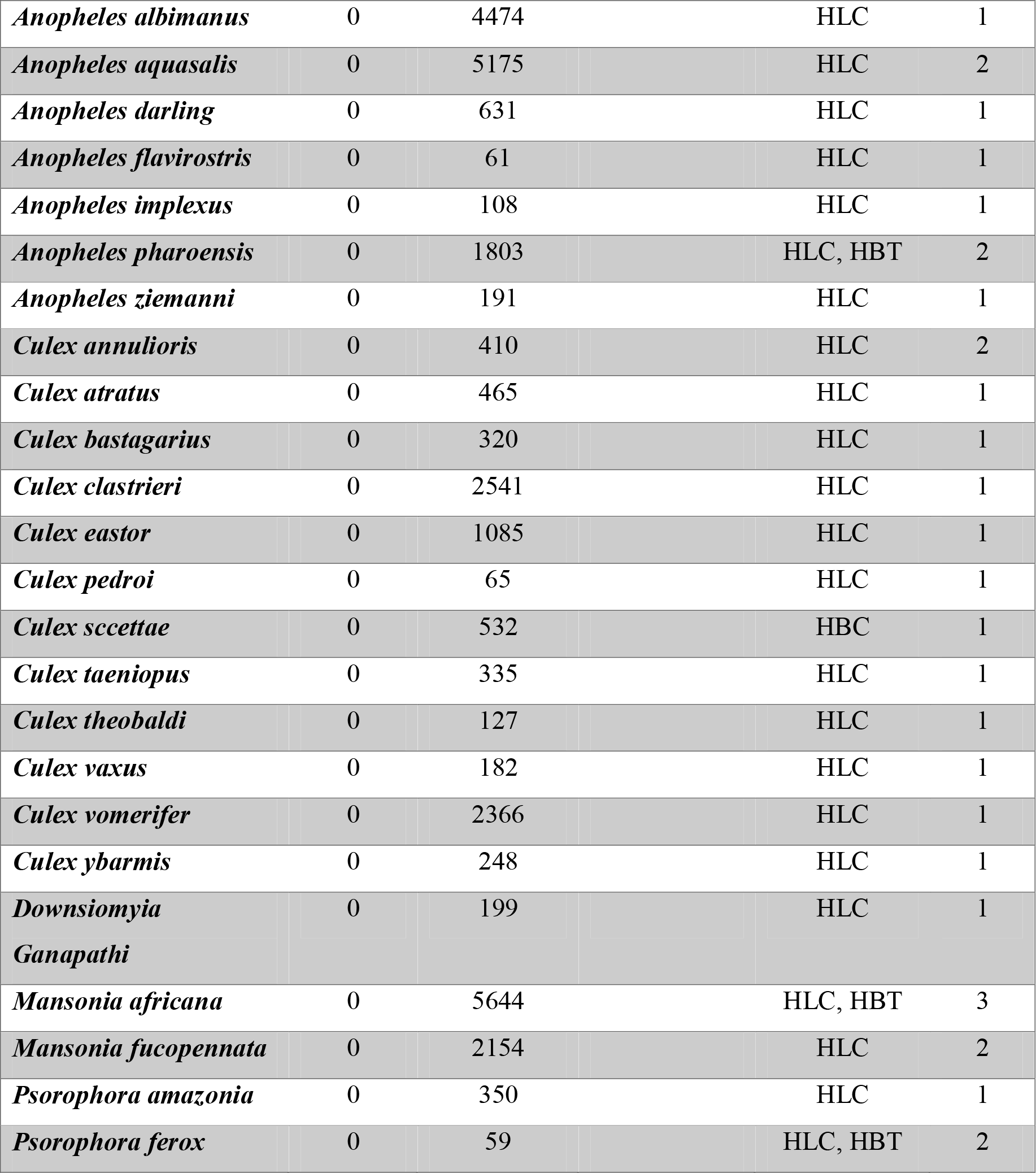
Numbers of females and males collected across mosquito species from the literature review of human-baited field collections. HLC = Human landing catch, HBT = Human baited trap, HBC = Human baited collection. We only present data for species with catches n > 50 individuals. Average proportion males was calculated by determining the proportion of males out of the total catch for each study, then averaging this proportion across studies. See table S4 for the complete dataset which includes proportions for each individual study.

| Species | Total males collected | Total females collected | Average proportion males | Collection method | # studies |
| --- | --- | --- | --- | --- | --- |
| **Species with male catches** |  |  |  |  |  |
| *Aedes aegypti* | 5880 | 12001 | 0.36 | HLC, HBT | 21 |
| *Aedes albopictus* | 7548 | 19667 | 0.19 | HLC, HBT, HBC | 19 |
| *Aedes flavipennis* | 14 | 39 | 0.26 | HLC | 1 |
| *Aedes furcifer* | 2 | 1987 | 0.05 | HLC | 2 |
| *Aedes polynesiensis* | 43 | 9225 | 0.005 | HBC | 1 |
| *Aedes riversi* | 90 | 125 | 0.42 | HBC | 1 |
| *Anopheles fuenstus* | 278 | 3703 | 0.025 | HLC, HBT | 3 |
| *Anopheles gambiae* | 115 | 15819 | 0.01 | HLC, HBT | 5 |
| *Armigeres malayi* | 2 | 869 | 0.002 | HLC | 1 |
| *Culex quinquefasciatus* | 711 | 1267 | 0.23 | HLC, HBT | 4 |
| *Culex riberensis* | 1 | 226 | 0.004 | HBC | 1 |
| *Culex vishnui* | 3 | 2983 | 0.001 | HLC | 1 |
| *Mansonia perturbans* | 246 | 12056 | 0.02 | HLC, HBT | 1 |
| *Mansonia uniformis* | 3 | 1747 | 0.03 | HLC | 2 |
| *Taeniorhynchus africanus* | 17 | 185 | 0.06 | HBT | 1 |
| *Taeniorhynchus uniformis* | 144 | 2418 | 0.06 | HBT | 1 |
| **Species without male catches** |  |  |  |  |  |
| *Aedes africanus* | 0 | 98 |  | HLC | 1 |
| *Aedes poicilius* | 0 | 125 |  | HLC | 1 |
| *Aedes serratus* | 0 | 125 |  | HLC | 1 |

|  |  |  |  |  |
| --- | --- | --- | --- | --- |
| *Anopheles albimanus* | 0 | 4474 | HLC | 1 |
| *Anopheles aquasalis* | 0 | 5175 | HLC | 2 |
| *Anopheles darling* | 0 | 631 | HLC | 1 |
| *Anopheles flavirostris* | 0 | 61 | HLC | 1 |
| *Anopheles implexus* | 0 | 108 | HLC | 1 |
| *Anopheles pharoensis* | 0 | 1803 | HLC, HBT | 2 |
| *Anopheles ziemanni* | 0 | 191 | HLC | 1 |
| *Culex annulioris* | 0 | 410 | HLC | 2 |
| *Culex atratus* | 0 | 465 | HLC | 1 |
| *Culex bastagarius* | 0 | 320 | HLC | 1 |
| *Culex clastrieri* | 0 | 2541 | HLC | 1 |
| *Culex eastor* | 0 | 1085 | HLC | 1 |
| *Culex pedroi* | 0 | 65 | HLC | 1 |
| *Culex scetetae* | 0 | 532 | HBC | 1 |
| *Culex taeniopus* | 0 | 335 | HLC | 1 |
| *Culex theobaldi* | 0 | 127 | HLC | 1 |
| *Culex vaxus* | 0 | 182 | HLC | 1 |
| *Culex vomerifer* | 0 | 2366 | HLC | 1 |
| *Culex ybarmis* | 0 | 248 | HLC | 1 |
| *Downsiomyia*<br>*Ganapathi* | 0 | 199 | HLC | 1 |
| *Mansonia africana* | 0 | 5644 | HLC, HBT | 3 |
| *Mansonia fucopennata* | 0 | 2154 | HLC | 2 |
| *Psorophora amazonia* | 0 | 350 | HLC | 1 |
| *Psorophora ferox* | 0 | 59 | HLC, HBT | 2 |

### 3.2. Human baited field collections

We conducted human-baited field trials in Australia in both temperate and tropical regions between 2014 and 2022. Over this period, we collected 13 mosquito species as shown in Table 2.

**Table 2.** Summary of human-baited field collections targeting male mosquitoes in Australia. Detailed information about collections can be found in Table S5. Köppen climate-zone codes: Am = Tropical Monsoon; Cfa = Humid subtropical; Cfb = Marina west coast; Csb: Mediterranean.

| Species | Males collected | Females collected | State | Köppen climate-zone |
| --- | --- | --- | --- | --- |
| *Aedes aegypti* | 89 | 140 | QLD | Am |
| *Aedes notoscriptus* | 0 | 441 | NSW | Cfa |
|  | 0 | 1501 | ACT | Cfb |
|  | 0 | 223 | VIC | Cfb |
|  | 0 | 1 | SA | Csb |
|  | 0 | 94 | QLD | Am |
| *Aedes alboannulatus* | 0 | 5 | ACT | Cfb |
|  | 0 | 5 | NSW | Cfa |
|  | 0 | 1 | VIC | Cfb |
| *Aedes vigilax* | 0 | 19 | NSW | Cfa |
|  | 0 | 2 | QLD | Am |
| *Aedes vittiger* | 0 | 3 | ACT | Cfb |
|  | 0 | 3 | QLD | Am |
| *Aedes procax* | 0 | 16 | NSW | Cfa |
| *Aedes rubrithorax* | 0 | 1 | ACT | Cfb |
|  | 0 | 21 | VIC | Cfb |
| *Culex orbostiensis* | 0 | 2 | NSW | Cfa |
|  | 0 | 5 | QLD | Am |
| *Culex quinquefasciatus* | 5 | 11 | VIC | Cfb |
| *Culex molestus* | 0 | 2 | VIC | Cfb |
|  | 0 | 9 | NSW | Cfa |
| *Culex annulirostris* | 0 | 5 | ACT | Cfb |
|  | 0 | 34 | NSW | Cfa |
|  | 2 | 38 | VIC | Cfb |
|  | 1 | 24 | VIC | Cfb |
|  | 0 | 1 | SA | Csb |
|  | 0 | 2 | ACT | Cfb |
| *Anopheles annulipes* | 1 | 1 | VIC | Cfb |
|  | 0 | 2 | ACT | Cfb |
| *Coquillettidia lienalis* | 0 | 2 | ACT | Cfb |

We found evidence of male attraction of Ae. aegypti to humans in our field collections, with males collected in 16/22 catches that captured this species. We also collected males from three other species (Cx. quinquefasciatus, Cx. annulirostris, and An. annulipes) but overall numbers were low. Aedes notoscriptus was by far the most prevalent mosquito captured, but no male individuals were collected despite recording thousands of females of this species.

### 3.3. Species-specific attraction of male Aedes mosquitoes to humans under laboratory conditions

In human-baited tent trials, we found no convincing evidence of attraction to humans in male Ae. notoscriptus or Ae. vigilax, with 5 or fewer observations of mosquitoes across all trials in each of the human-baited and unbaited treatments (Figure 2). Males of both species were inactive in the presence of human subjects and typically rested on the walls of the tent. However, we observed consistent attraction to humans in Ae. aegypti (Figure 2). The number of males observed in human-baited treatments after 30 min was significantly higher than in unbaited treatments across all tested human subjects (Wilcoxon signed-rank test: landed: z = 1.072, p < 0.001; in flight: z = 7.755, p < 0.001; total: z = 1.056, p < 0.001). Attraction was persistent, with males observed in flight around human subjects for the entire 30 minutes.

**Figure 2.**
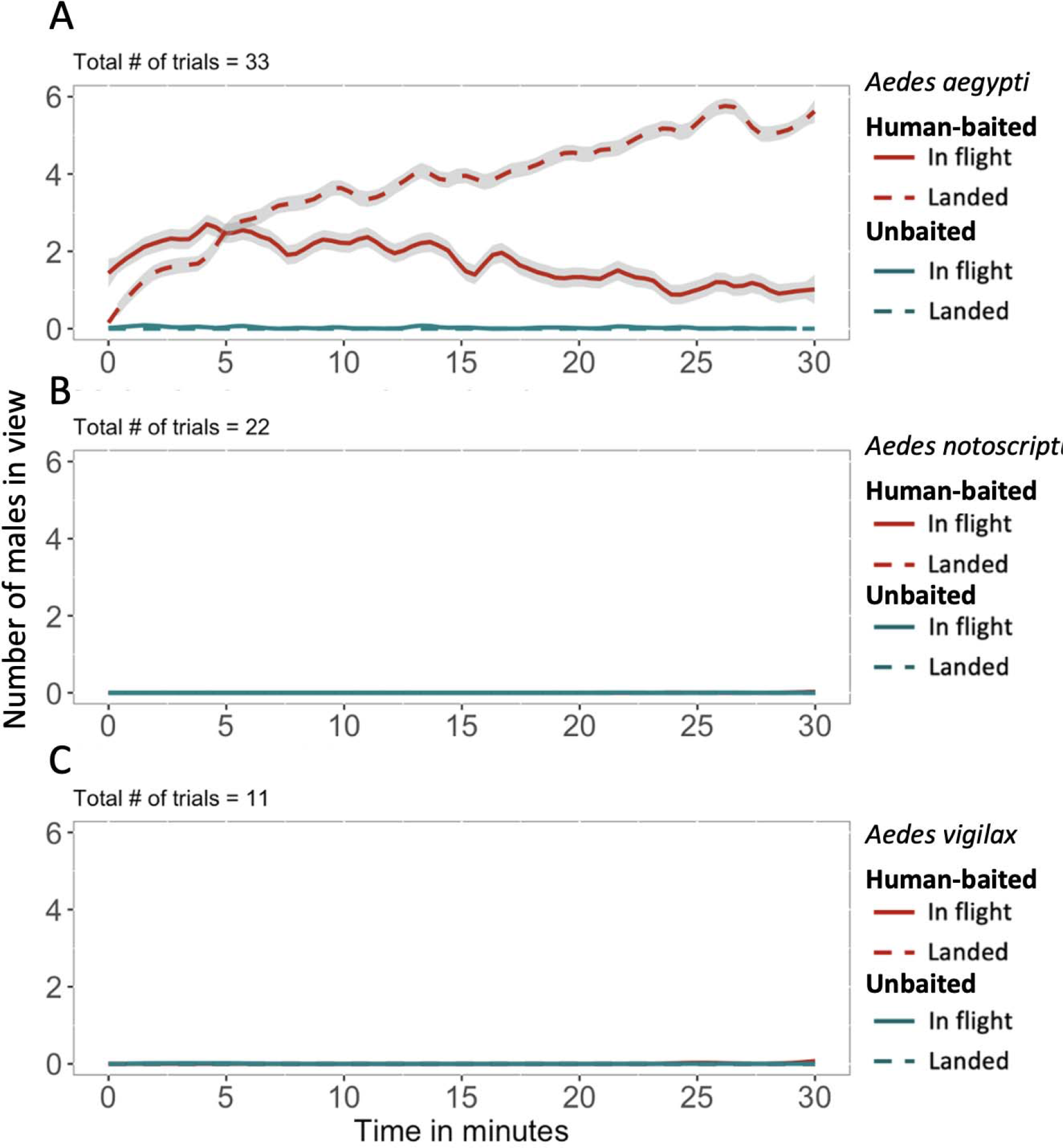
Comparison of male attraction to humans for three *Aedes* species in tent trials. The number of male mosquitoes of *Aedes aegypti* **(A)**, *Aedes notoscriptus* **(B)** and *Aedes vigilax* **(C)** observed in view of a camera every 20 s. Mosquitoes that were in-flight and landed are shown with solid and dashed lines respectively. Human-baited treatments are indicated in red, with unbaited controls shown in blue. 95% confidence intervals are shown in grey. Data were averaged across all human subjects, with data for *Ae. aegypti* males presented separately for each human volunteer in Figure S3.

Additionally, we observed an increasing number of males that had landed on the subject throughout the trials (Figure 2). While human subjects were not compared directly in this experiment, mosquito observations were much higher for some subjects, suggesting differential attraction (Figure S3).

### 3.4 Mosquito preferences for different human subjects

We observed significant host preferences among male and female Ae. aegypti and female Ae. notoscriptus in pairwise comparisons between five human subjects (ANOVA: Ae. aegypti males: F = 5.019, df = 4, 15, p = 0.008; Ae. aegypti females: F = 5.81, df = 4, 15, p = 0.005; Ae notoscriptus females: F = 4.137, df = 4, 15, p = 0.018). Although less pronounced, male Ae. aegypti showed a preference for the same human subjects as female Ae. aegypti (Figure 3). Tukey’s posthoc tests showed that significantly more mosquitoes were attracted to certain human subjects over others (Ae. aegypti males: Subject A vs Subject E: p = 0.009; Subject B vs Subject E: p = 0.017; Ae. aegypti females: Subject A vs Subject D: p = 0.025; Subject A vs Subject E: p = 0.04; Ae notoscriptus females: Subject B vs Subject D: p = 0.035; Subject B vs. Subject E: p = 0.03) (Figure 3). We found similar results when analysing the original data prior to calculation of indices (GLMER: Ae. aegypti males: df = 157, p < 0.001; Ae. aegypti females: df = 157, p < 0.001; Ae. notoscriptus females: df = 119, p < 0.001). Tukey’s posthoc tests showed that significantly more mosquitoes were attracted to certain human subjects over others (Ae. aegypti males: Subject A vs Subject E: p = 0.039; Ae. aegypti females: Subject A vs Subject D: p = 0.039; Subject A vs Subject E: p = 0.045; Ae notoscriptus females: Subject B vs Subject D: p = 0.047; Subject B vs. Subject E: p = 0.041).

**Figure 3.**
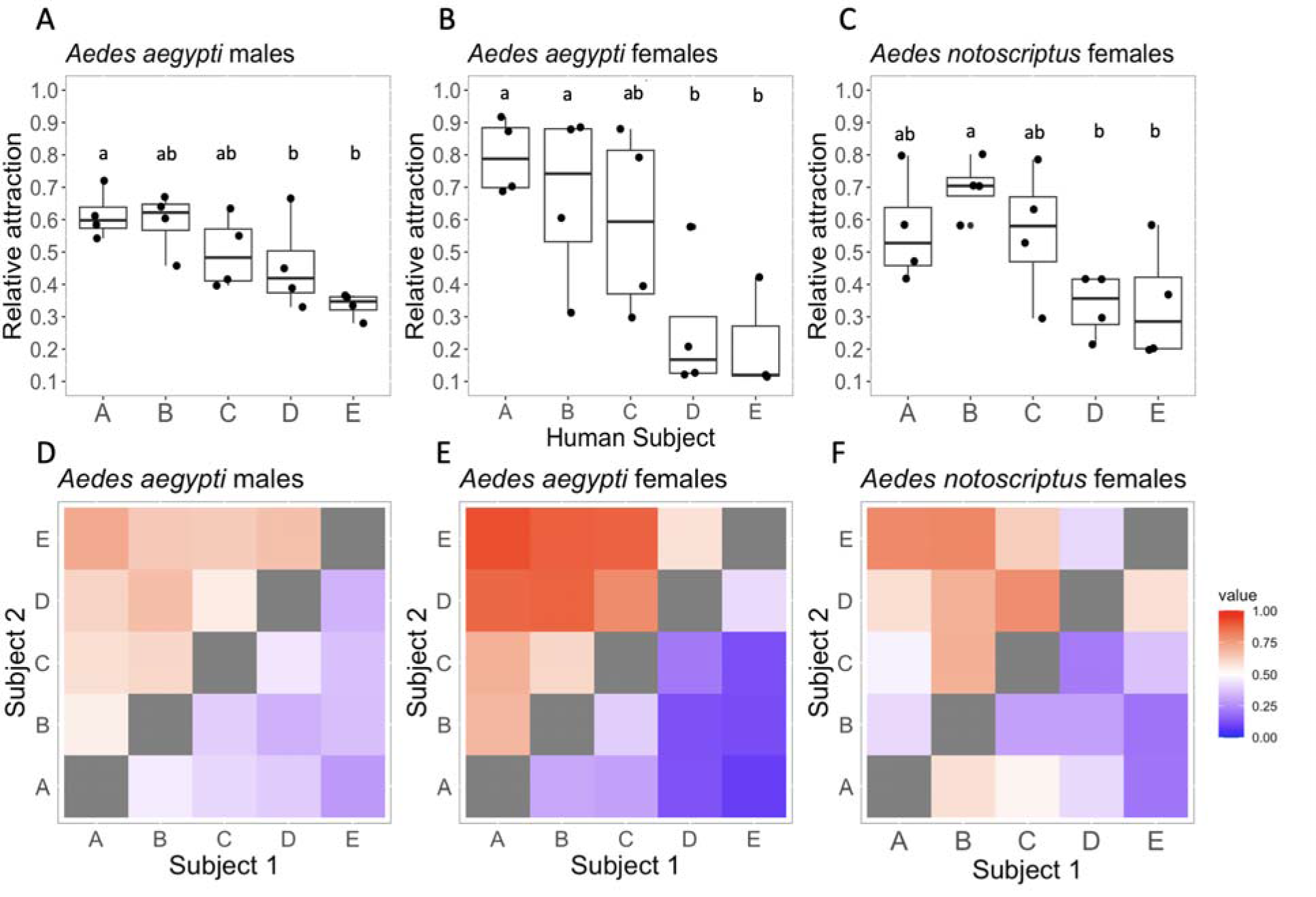
Relative attraction of female *Aedes notoscriptus* and male and female *Aedes aegypti* to different human subjects in pairwise comparisons. The upper row **(A-C)** shows boxplots of relative attraction between the five human subjects across *Ae. aegypti* males **(A)** and females **(B)** and *Ae. notoscriptus* males **(C).** The preference index was calculated by dividing the number of mosquitoes attracted to one human subject over the number of mosquitoes attracted to both subjects. Dots represent the mean attraction of the relevant subject to the other four subjects across 8 replicate trials. Comparisons with significant (P < 0.05) pairwise differences are indicated by different letters. The lower row **(D-F)** presents heat maps displaying the preference index in pairwise comparisons between human subjects. Preference indices are shown on a 0-1 scale, with higher values (red) indicating stronger attraction to subject 1, lower values (blue) indicating stronger attraction to subject 2, and 0.5 (white) indicating no preferential attraction between pairs of human subjects.

Jonckheere-Terpstra tests comparing preference index values for the focus subject against the other subjects ranked in order of overall attractiveness showed that the attractiveness to one subject was not influenced by the other human subject present in the pairwise comparison; this lack of dependence on the other subject was found for Ae. aegypti females and males as well as for Ae. notoscriptus females (Table S3). Mantel tests on preference index matrices between Ae. aegypti males and females as well as Ae. notoscriptus females were positive but not significant (Table S3), suggesting a similar pattern of preferences for human subjects among the three groups.

### 3.5 Effect of mosquito repellent on male mosquitoes that show attraction to humans

Commercial mosquito repellent applied to exposed skin was effective in reducing the attraction of male Ae. aegypti to human subjects (Figure 4). Significantly fewer mosquitoes landed on the exposed skin of human subjects wearing repellent (t-test: t = 6.51, df = 8, p < 0.001). Furthermore, fewer male Ae. aegypti mosquitoes were observed flying in field of view of the camera towards humans wearing repellent compared to untreated subjects (t-test: t = 8.18, df = 8, p < 0.001).

**Figure 4.**
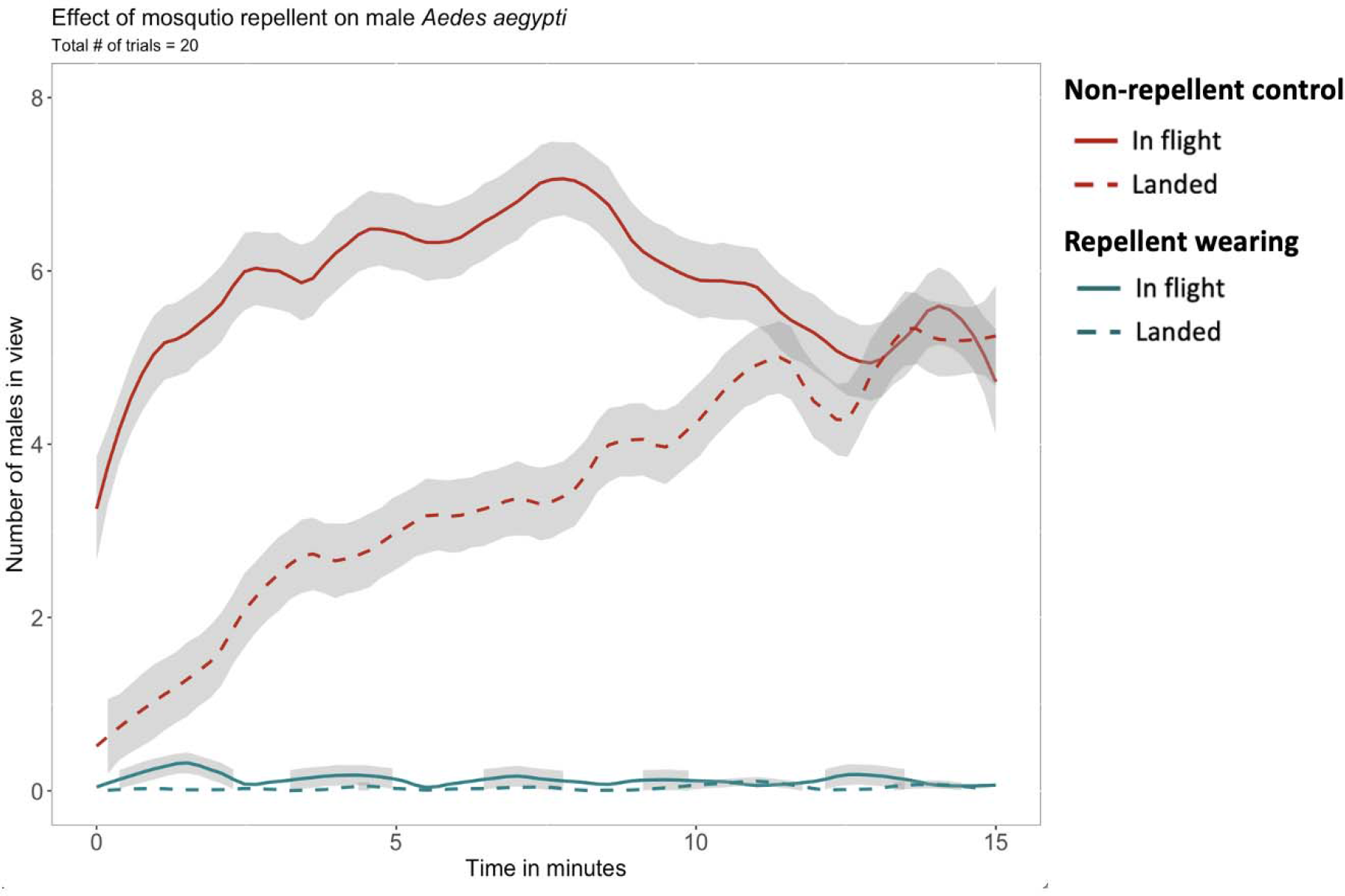
Effect of mosquito repellent applied to exposed skin on swarming and landing by male *Aedes aegypti*. The number of male *Ae. aegypti* in view of a camera was recorded every 20 s. Mosquitoes that were in-flight and landed are shown with solid and dashed lines respectively. Repellent-wearing treatments are indicated in red, with non-repellent controls shown in blue. 95% confidence intervals are shown in grey. Data were pooled across all human subjects.

## Discussion

In this study we presented an integrated approach that combines a literature review with our own field collections and laboratory experiments to investigate the phenomenon of male mosquito attraction to humans. The literature review indicated that male attraction to humans is apparent in only a limited number of species, including Ae. aegypti and Ae. albopictus. Our human-baited field collections were consistent with the review, where we observed clear evidence for attraction to humans in male Ae. aegypti only among the 13 captured mosquito species. Subsequently, in laboratory experiments, we assessed the attraction of male mosquitoes from different species and found that Ae. notoscriptus and Ae. vigilax males exhibited no discernible attraction to humans, whereas male Ae. aegypti consistently displayed attraction for the full duration of the trials. Remarkably, both male and female Ae. aegypti demonstrated similar preferences for different human subjects, suggesting that male Ae. aegypti respond to similar cues as their female counterparts. Additionally, we found that repellent not only reduces landing of male mosquitoes on humans but also decreases swarming behaviour. Even though males do not bite, they can still be regarded as a nuisance, as reported in some communities (https://www.todayonline.com/voices/project-wolbachia-residents-are-killing-helpful-mosquitoes-which-can-be-nuisance).

Our literature review revealed a scarcity of observational data on male mosquito attraction to humans. This is primarily due to the limited reporting of male catches in field studies specifically designed to capture or assess their attraction (e.g., 39–41). The diverse nature of the results made a traditional meta-analysis inappropriate, leading us to classify our approach as a literature review. Human landing catches (HLC) were commonly employed for mosquito collection in the field (46), but they can introduce bias by collecting more females than males. Males, even if attracted to humans, often fly around without landing, resulting in a higher collection rate of females. Furthermore, variations in HLC execution across studies make it challenging to ensure comparability of results. Most of the screened studies primarily focused on collecting female mosquitoes or testing female attraction to different traps or hosts, with limited consideration given to males or reporting of male catch data. Additionally, many studies lacked clear information on whether reported catch numbers encompassed all observed species or only those relevant to the study, potentially leading to under-sampling of certain species. This lack of clarity may contribute to an overrepresentation of species like Ae. aegypti and Ae. albopictus, which are commonly recognized as nuisance or vector species. Therefore, caution should be exercised when interpreting the findings of our literature review. Despite these limitations, we identified a distinct pattern in male attraction to humans, with highly anthropophilic and invasive species (e.g., Ae. aegypti, Ae. albopictus, Cx. quinquefasciatus) displaying greater attraction compared to species with broader host preferences and lower invasiveness (Figure 1, Table 1). Our own field collections targeting males support these findings, as we consistently observed male attraction in Ae. aegypti, while other species showed either minimal or no male attraction (Table 2).

Our laboratory experiments comparing different Aedes species provide clear evidence that male attraction to humans is a species-specific phenomenon. Male Ae. aegypti persistently swarmed and landed on humans, while Ae. notoscriptus and Ae. vigilax displayed no attraction (Figure 2). Our results also indicate that male Ae. aegypti exhibit varying levels of attraction towards different human participants (Figure 3; Figure S3), a phenomenon well documented in female mosquitoes of different species [41,42], including Ae. aegypti [15,37,38,43,44]. Consistent preferences for specific human subjects were found across females of Ae. aegypti and Ae. notoscriptus, indicating that these species respond to similar host-specific cues. These findings are noteworthy as Ae. notoscriptus feeds on a broader range of hosts [45] compared to Ae. aegypti, and it is important to acknowledge that blood feeding patterns may not necessarily reflect host preferences as they could also be influenced by host availability [46].

Male Ae. aegypti demonstrated similar individual host preferences as female Ae. aegypti (Figure 3). The attraction of mosquitoes to humans is a complex process that depends on multiple cues being identified and integrated even at long distances [47]. Our data suggest that components of this process may be similar across males and females. Mosquito genome studies have identified several receptor families that detect volatile chemicals [48–50]. Studies investigating the Ae. gambiae ionototropic receptor family have revealed that the expression of receptors was largely similar between the sexes, but males generally have a lower expression level of all receptors [51], suggesting that they may be responsive to the same chemical compounds as females, but at a reduced sensitivity. Amos et al [34] described long-range attraction of Ae. aegypti males but no detectable short-range attraction, suggesting that males can integrate multiple cues associated with humans for long distance attraction, but sexes respond differently to close distance cues which can be different to cues required for long distance integration [47]. At close distances males may respond to different cues (e.g., room for swarming). Recent research has revealed that preferences in female Ae. aegypti for specific humans is influenced by their skin-derived carboxylic acid levels [38], and males may also detect this odour cue since they show a similar preference to different humans in our experiments (Figure 3). While our results show a similar preference pattern between male and female Ae. aegypti, it is important to note that male and female attraction were measured in different ways (tent trials vs. a two-port olfactometer) which could have introduced differences in overall preference levels.

The species-specific attraction to humans shown by male mosquitoes raises intriguing questions about the evolution of this behavioural variation. Males of several species, including Ae. albopictus and Ae. aegypti aggregate in swarms near hosts in nature [27,29,30,52]. Females entering these swarms are engaged by males, leading to copulation [30]. Both species are active and bite during the day, which might lead to host seeking and mating behaviour being coupled processes [52,53]. Males of Ma. uniformis and Ma. africana also reportedly orient towards non-human animals in search of females for mating [54,55]. In Anopheline and Culicine mosquitoes, swarming behaviour does not require hosts [56,57]. Anopheles gambiae form large swarms in the absence of host animals, likely relying on visual cues [57]. This species exhibits nocturnal feeding and crepuscular mating patterns, and the separation of feeding and mating at different times may factor into the lack of male attraction to hosts. Males of other species may target different habitats for mating; for instance, Ae. polynesiensis mates near larval habitats and exhibits higher insemination rates there than Ae. aegypti [58]. In species such as Ae. communis and Ae. stimulans, swarming is a pre-requisite for mating and has been observed in large walk-in cages, with mating pairs forming in flight [59]. These observations point to a diversity of mating strategies and help explain the lack of males collected for many of the species in our literature review and field collections.

Developing an understanding of male mating behaviour is important because successful mating with wild females is critical for mass-reared male mosquitoes released for disease control efforts [60–62]. However, being able to facilitate the right circumstances for this when planning releases is a challenge without knowing the factors that influence mating behaviour. Male mosquito release programs need to consider what species-specific mating and host-seeking behaviour their target species displays. For example, releases with mosquitoes including Ae. aegypti should consider that the presence of humans may be important for inducing mating, while releases of An. gambiae should focus on other factors and areas away from humans that induce this behaviour. Finding the right species-specific swarming marker or cues will be useful for the development of efficient male trap techniques to benefit surveillance.

Mating behaviour is also important in the establishment and maintenance of laboratory colonies. For example, Watson et al [63] argued that difficulties to establish Ae. notoscriptus colonies in the laboratory stems from mating behaviour that cannot easily be facilitated in cages. Understanding these behaviours can help researchers to identify the best methods for maintaining colonies, such as using bigger cages with larger numbers of males to induce swarming, adding swarm markers such as plants or providing host odours if the species shows male attraction to hosts. Furthermore, understanding the mating behaviour of mosquitoes can help researchers to investigate the evolution of different mating strategies and how they influence the population dynamics of mosquitoes, as well as the underlying genetic and physiological mechanisms that drive these behaviours.

## Conclusion

In conclusion, our study presented a comprehensive examination of male mosquito attraction to human hosts through a combination of literature review, field collections, and laboratory experiments. We demonstrated species-specific attraction patterns, with male Ae. aegypti showing persistent attraction and landing on humans, while other species such as Ae. notoscriptus and Ae. vigilax exhibited no significant attraction. The effectiveness of mosquito repellents on male mosquitoes attracted to humans was also evaluated, showing promising results in reducing landing and swarming behaviour. Further investigations are needed to explore the efficacy of repellents on male mosquitoes that have been sterilized using various methods, such as Wolbachia infection or exposure to x-rays. Additionally, our findings underscore the importance of understanding species-specific mating behaviour and its implications for mosquito control efforts, such as targeted release programs and laboratory colony maintenance. Further research on male mosquito attraction, mating behaviour, and the underlying genetic and physiological mechanisms will contribute to our knowledge of mosquito population dynamics and aid in the development of effective control strategies.

## Acknowledgements

The authors would like to thank Sophie Collier, Jessica Home, Ashritha Dorai, Eloïse Ansermin, Courtney Brown, Sian McDonald, Nick Bell, Xinyue Gu, Ella Yeatman, Sonia Sharma, John Black, Nancy Endersby-Harshman, Jake Brown, Allister Small, and Xuefen Xu for their help with experiments and field collections. We also thank Brendan Trewin for providing Ae. vigilax eggs, as well as Leon Hugo who provided Ae. notoscriptus eggs. This research was supported by the Robert Johanson and Anne Swann Fund (to PAR) as part of the Big Science Pitch at the University of Melbourne. VP was financially supported by the Australian Government Research Training Program Scholarship. PAR was supported from Aalborg University to AAH. AAH was funded by the National Health and Medical Research Council Partnership Project grant 1196396 ‘Stopping Buruli ulcer in Victoria’.

## Data accessibility

Data can be accessed here: https://datadryad.org/stash/share/ypDPmDRfv1kkG4Nhc2VROhT-V35Pl9RLAkX4C6eLRfE(Data accessible after publication here: https://doi.org/10.5061/dryad.tb2rbp04x)

