## Supplementary material for "How often are male mosquitoes attracted to humans?": Figures S1 and S2

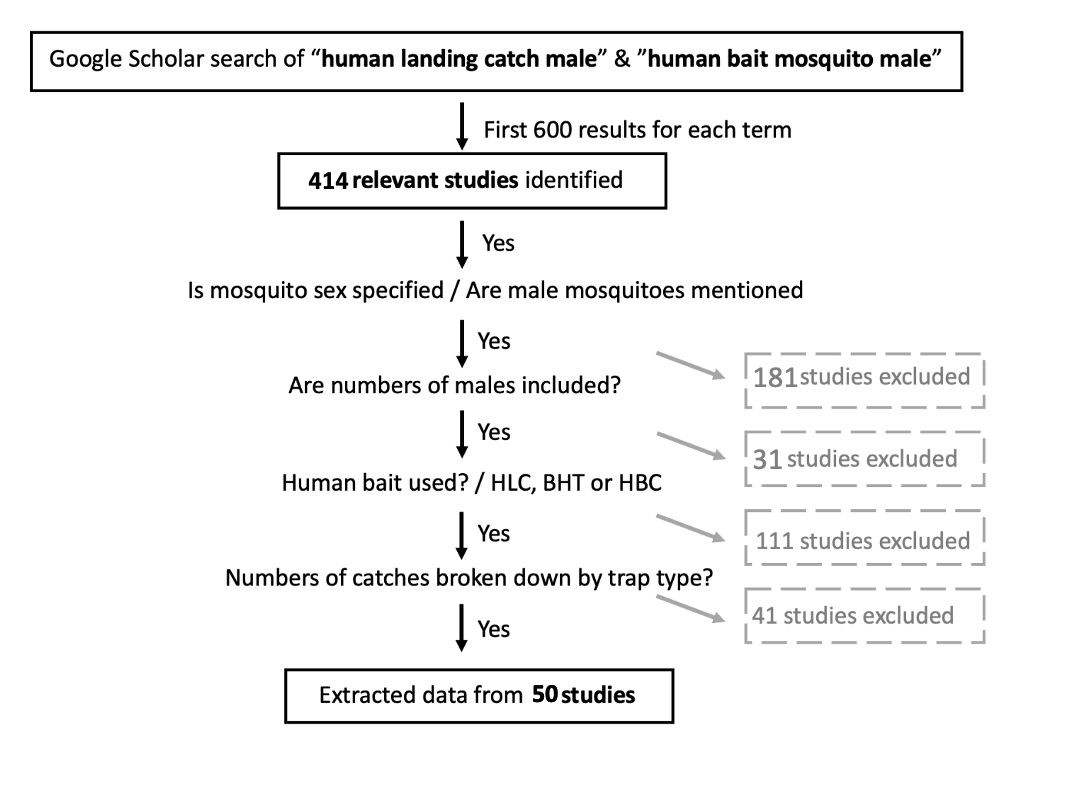

**Figure S1** Outline of inclusion criteria of the literature review presented in this study.

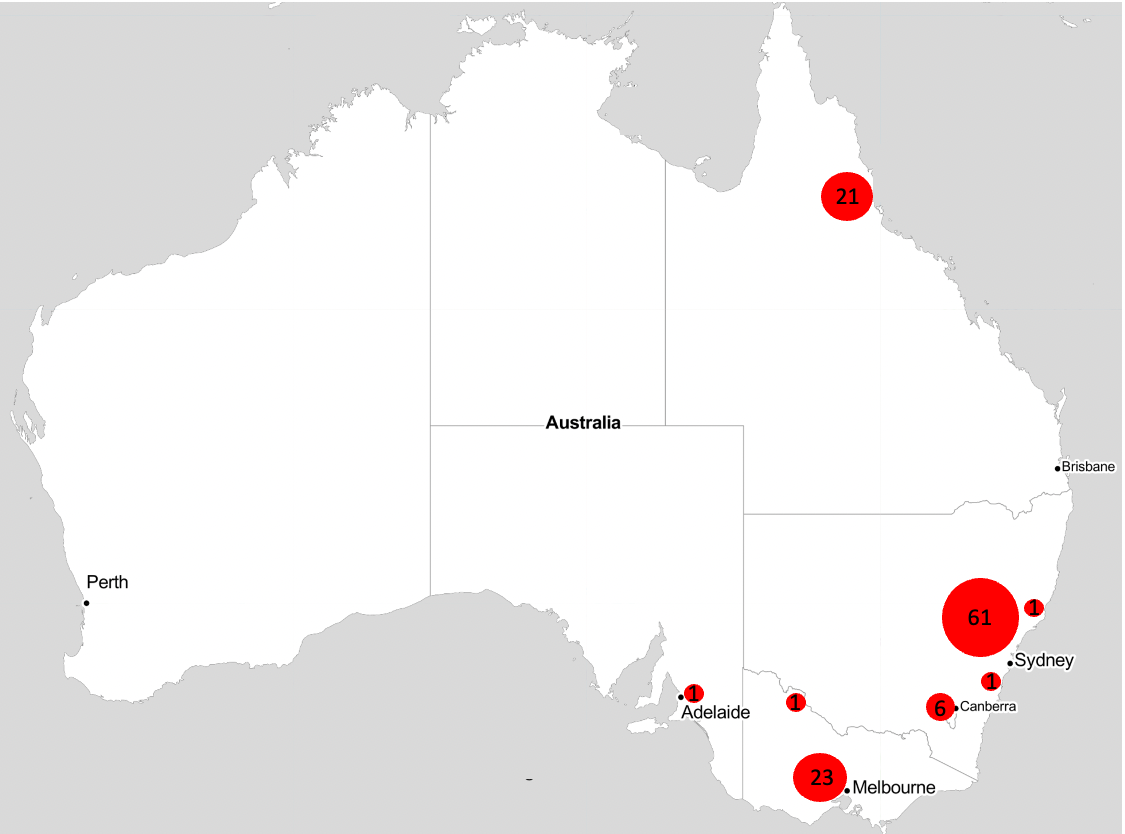

**Figure S2** Number of human-baited field collections across VIC, SA, ACT, NSW and QLD. The size of the circles indicates the number of collections, as well as numbers in circles.

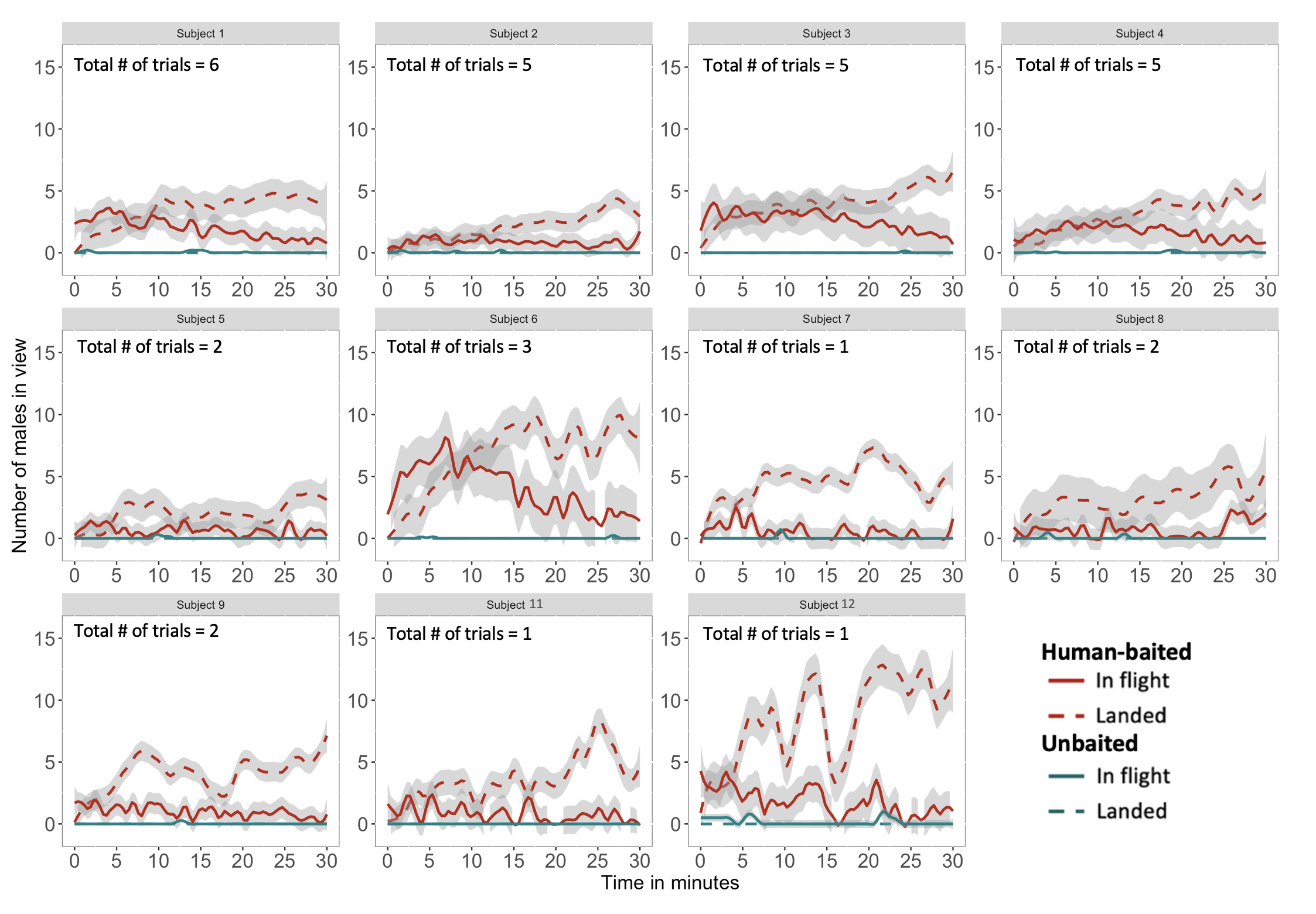

**Figure S3. Comparison of male *Aedes aegypti* attraction to humans split by human subjects.** Presented is the number of male mosquitoes of *Aedes aegypti* observed in view of the camera every 20 s. Mosquitoes that were in-flight and landed are shown with solid and dashed lines respectively. Human-baited treatments are indicated in red, with unbaited controls shown in blue. 95% confidence intervals are shown in grey.

**Table S1.** Number of trials, human subjects and days of the tent experiments across species.

| **Species** | **# trials** | **# human subjects** | **# days** |
| --- | --- | --- | --- |
| ***Ae. aegypti*** | 33 | 11 | 6 |
| ***Ae. notoscriptus*** | 22 | 8 | 4 |
| ***Ae. vigilax*** | 11 | 7 | 2 |

**Table S2.** Mantel tests on preference index matrices between *Ae. aegypti* females and males and *Ae. notoscriptus* females.

| **Species / sex Group 1 vs Species / sex Group 2** | | **P-value** | **Mantel r** |
| --- | --- | --- | --- |
| *Aedes aegypti* males | *Aedes aegypti* females | 0.1038 | 0.383 |
| *Aedes aegypti* males | *Aedes notoscriptus* females | 0.1783 | 0.326 |
| *Aedes aegypti* females | *Aedes notoscriptus* females | 0.0950 | 0.620 |

**Table S3.** Jonckheere-Terpstra tests on preference index data for *Ae. aegypti* females and males and for *Ae. notoscriptus* females where the attractiveness of each human subject was first ranked, and the preference of each human subject was tested against the other 4 subjects ordered in terms of their attractiveness using data from the 5 replicates. The test statistic (JT) measures the strength of evidence against the null hypothesis that there is no trend in the data.

| ***Ae. aegypti* females** | | |
| --- | --- | --- |
| **Human subject** | **Jonckheere – Terpstra statistic** | **P-value** |
| A | 6 | 0.0652 |
| B | 9 | 0.2821 |
| C | 10 | 0.3922 |
| D | 9 | 0.2822 |
| E | 6 | 0.0592 |
| ***Ae. aegypti* males** | | |
| A | 6 | 0.0606 |
| B | 9 | 0.2748 |
| C | 10 | 0.3992 |
| D | 9 | 0.2850 |
| E | 6 | 0.0636 |
| ***Ae. notoscriptus* females** | | |
| A | 10 | 0.392 |
| B | 10 | 0.3844 |
| C | 11 | 0.5619 |
| D | 10 | 0.386 |
| E | 7 | 0.1116 |
